# Long-term genomic coevolution of host-parasite interaction in the natural environment

**DOI:** 10.1101/101576

**Authors:** Elina Laanto, Ville Hoikkala, Janne Ravantti, Lotta-Riina Sundberg

## Abstract

The antagonistic coevolution of parasite infectivity and host resistance alters the biological functionality of species, with effects spanning to communities and ecosystems. Still, studies describing long-term host-parasite coevolutionary dynamics in nature are largely missing. Furthermore, the role of host resistance mechanisms for parasite evolution is poorly understood, necessitating for the molecular and phenotypic characterization of both coevolving parasites and their hosts. We combined long-term field sampling (2007-2014), *in vitro* cross-infections and time-shift experiments with bacteriophage whole genome sequencing and bacterial (*Flavobacterium columnare*) CRISPR (Clustered Regularly Interspaced Short Palindromic Repeats) profiling to show the molecular details of the phage-bacterium arms race in the environment. Bacteria were generally resistant to phages from the past and susceptible to phages in the future. The bacterial resistance selected for increased phage infectivity and host range, correlating directly with the expansion of phage genome size by 2656 bp. In the bacterial host, two CRISPR loci were identified: a type II-C locus and an RNA-targeting type VI-B locus. While maintaining a core set of conserved spacers, phage-matching spacers appeared in the variable end of both CRISPR loci over time. The appearance of these CRISPR spacers in the bacterial host often corresponded with arms race -manner molecular changes in the protospacers of the coevolving phage population. However, the phenotypic data indicated that the relative role of constitutive defence may be more important in high phage pressure, highlighting the importance of our findings for understanding microbial community ecology and in the development of phage therapy applications.

## Introduction

One of the fundamental questions in evolutionary biology and in medicine is how host immunity drives the evolution of pathogens, as all living organisms are exposed to infections that can cause significant harm and selection in host population. The antagonistic coevolutionary arms race of parasite infectivity and host resistance leads to adaptations and counter-adaptations in the coevolving partners^1-4^, and also has a central role in the evolution of host-parasite relationships in the microbial world. This has been shown especially under experimental settings, where lethal infections by bacterial viruses, (bacterio) phages, shape the diversity and dynamics of host bacterial populations^1,2,4,5^, whereas the phages have the capacity to rapidly overcome host immunity^1,6-10^. It has been suggested, however, that the adaptive potential of phage and high costs in phage resistance in bacteria can limit the phage-bacterium coevolution^11^. Yet, the arms race dynamics observed in laboratory experiments substantially differ from the real-life dynamics under the complex web of surrounding interactions present in the environment, which impacts the ecology and evolution of both phages and their host^12,13^.

CRISPR (Clustered Regularly Interspaced Short Palindromic Repeats) and associated cas genes form the bacterial CRISPR/Cas adaptive immune system against phages and plasmids. CRISPR/Cas protects bacteria from infections by cleaving invading nucleic acid sequences (protospacers) that are identical or nearly identical to the spacers in the bacterial CRISPR repeat-spacer array^14,15^. Protospacer adjacent motifs (PAMs) are used to distinguish self from non-self and are crucial in most CRISPR systems^16^ while protospacer flanking sites (PFSs) determine the efficiency of interference in RNA-targeting CRISPR systems, but have no autoimmunity related functions^17^. Over time, novel spacers accumulate in one end of the array and may therefore confer resistance to multiple phages. However, in accordance with antagonistic coevolution theory, phages can counter-adapt to host immunity via point mutations in the protospacer or PAM/PFS sequences^6,18,19^.

Following evolutionary change in natural communities is challenging, and linking genomic change data with the coevolutionary dynamics of phages and bacteria is one of the key challenges in microbiology. Therefore, a comprehensive view of host-parasite coevolutionary dynamics in the environment, surrounded by a network of other trophic interactions, is missing. In fact, previous studies have concentrated either on phenotypic changes^8,20^ or CRISPR genetics using metagenomic approaches^21-23^. Particularly, empirical evidence on the relative importance of constitutive and adaptive resistance (cell surface modifications and CRISPR loci insertions, respectively^24^) in bacteria and the corresponding changes in the parasitic phage^25^ supported by phenotypic data from natural settings is missing. Different resistance mechanisms are predicted to be important in different ecological conditions, therefore teasing apart the relative role of different bacterial resistance mechanisms is important for understanding microbial community ecology. From applied perspective, such information is also crucial for the development of long-term functional phage therapy applications^26^.

Here, we characterize the coevolution of populations of bacteriophages and their bacterial hosts (*i.e.* the fish pathogen *Flavobacterium columnare*^27^) at the phenotypic and genetic level, in a flow-through aquaculture setting during the period 2007-2014. We sampled for bacterial and phage isolates from the fish farming facilities or immediate surroundings and performed cross-infections to analyse bacterial resistance patterns. To link the phenotypic patterns with molecular changes, we sequenced phage genomes to understand the determinants of the host range and characterized the bacterial CRISPR loci to understand the role of adaptive immunity in driving genome evolution and host range in the phage population. We report the patterns of arms race coevolution in a natural community and demonstrate that whereas bacteria evolve resistance, phages evolve a broader host range over time, which is associated with increase in genome size. We also observed evolutionary change in phage genomes in response to bacterial adaptive (CRISPR) and constitutive (surface modification) immunity, exemplifying the importance of both resistance mechanisms in natural bacterial populations. In addition, in our knowledge, this is the first study to demonstrate the evolution of an RNA-targeting CRISPR system in a natural context.

## Results

### Phage and bacterial isolates

*F. columnare* phages infect the host in a genotype-specific manner^28^, which allowed us to analyze both phage and host isolates over long time scales in locally adapting populations at a fish farm in Central Finland. During the period 2007-2014, we isolated 17 *F. columnare* isolates that belong to the genetic group C (Supplementary Table S1) and 30 dsDNA phages (belonging to the family *Myoviridae*) infecting specifically this host group (Supplementary Table S2).

### Coevolution of phage infectivity and bacterial resistance over time

First, we analysed the phage host range by cross-infecting all 30 phage isolates with all 17 bacterial isolates. The most recently isolated phages had the widest host range, being able to infect nearly all bacterial hosts. Analysed using a time-shift approach^29^ the phage isolation time point had a significant effect on infectivity compared to the bacterial host. The bacteria were in general resistant against phages from the past but susceptible to infection by phages from contemporary and future time points (24% and 18% resistant, respectively) (F_(2, 507)_=17,464, p<0.001) (Figure 1a). The phage isolation year (included as a random factor in this analysis) had a significant effect (Wald Z=4.014, p<0.001). A trend in bacterial phage resistance was also observed when the age difference between phage and bacterial isolates was analysed by years (F_(9, 500)_=1.664, p=0.095, phage isolation year as a random factor: Wald Z=3.39, p=0.001, Figure 1b). All bacteria were resistant to phages from 3-4 years from the past, and, on the other hand, bacterial resistance was below 0.5% against phages from 4-5 years in the future.

**Figure 1.**
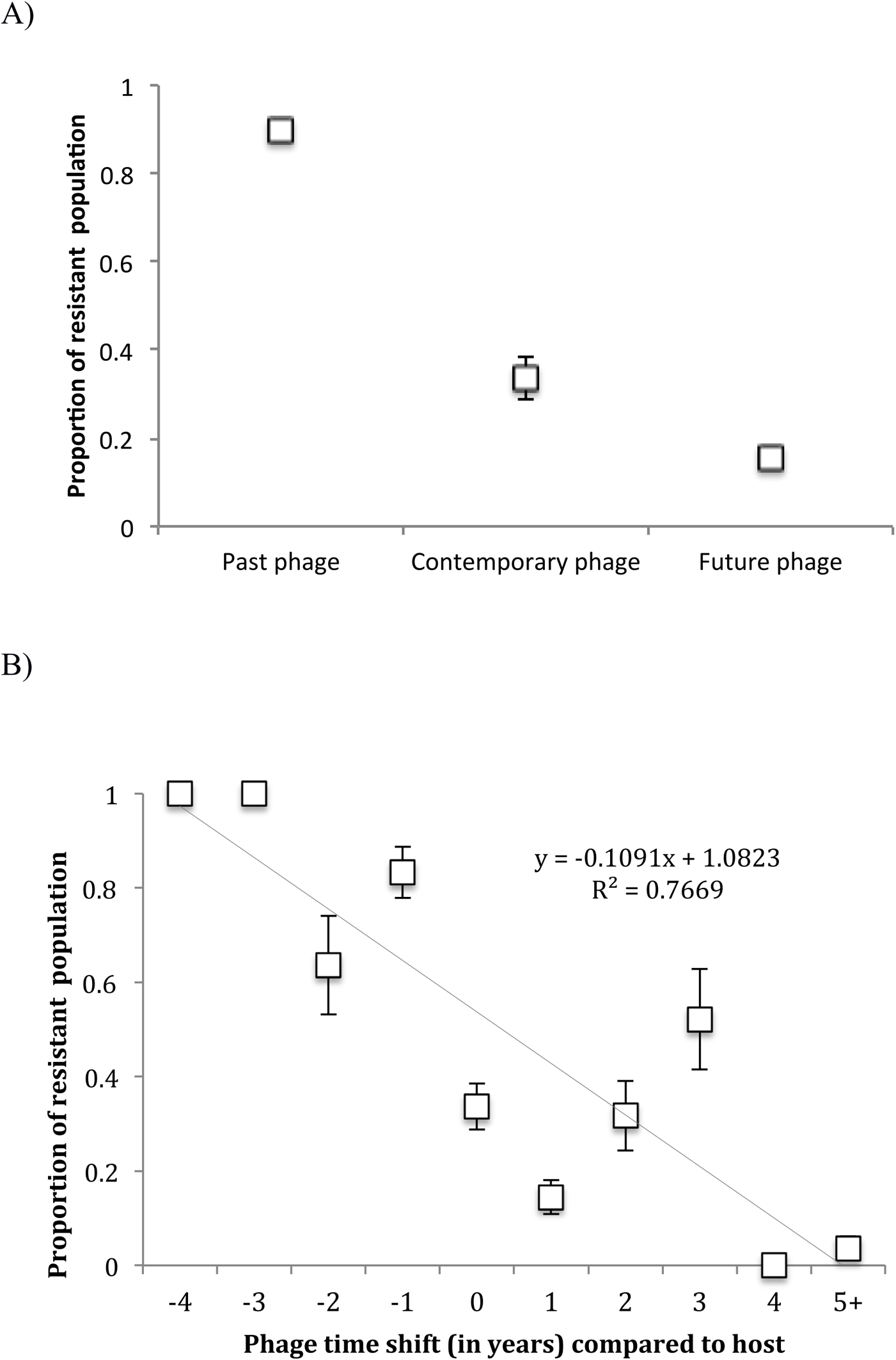
Time shift of phage-bacterium coevolution at fish farming environment measured as mean proportion (+/- S.E.) of resistant host population. A) Host resistance when exposed to phage from past, contemporary and future time points. B) Annual relative change in host resistance when exposed to phages from the past (−4 to −1 years), contemporary (0) and future (+1 to +5 years).

### Host resistance drives phage genome evolution

To understand phenotypic resistance at the genomic level, we sequenced genomes of 17 phage isolates from 2009 to 2014. This resulted in complete genomes with lengths ranging from 46 481 bp to 49 084 bp (Figure 2, Supplementary Table S3 and S4, Supplementary Figure S1). Phage genomes from different years were highly similar, enabling us to study differences at the nucleotide level. From the predicted 73 (2009) to 76 (2014) ORFs, 52 were 100% identical between all genomes, including putative terminase, portal protein and several structural proteins (e.g. major capsid protein). Differences in putative tail and structural proteins (two phages from 2011 and three from 2014, in ORFs 27, 28, 36 and 37, Supplementary Figure S1), suggest a conventional arms race between phage infectivity and host resistance via receptor mutation in natural settings (see^9,10^ for similar experimental evolution results).

**Figure 2.**
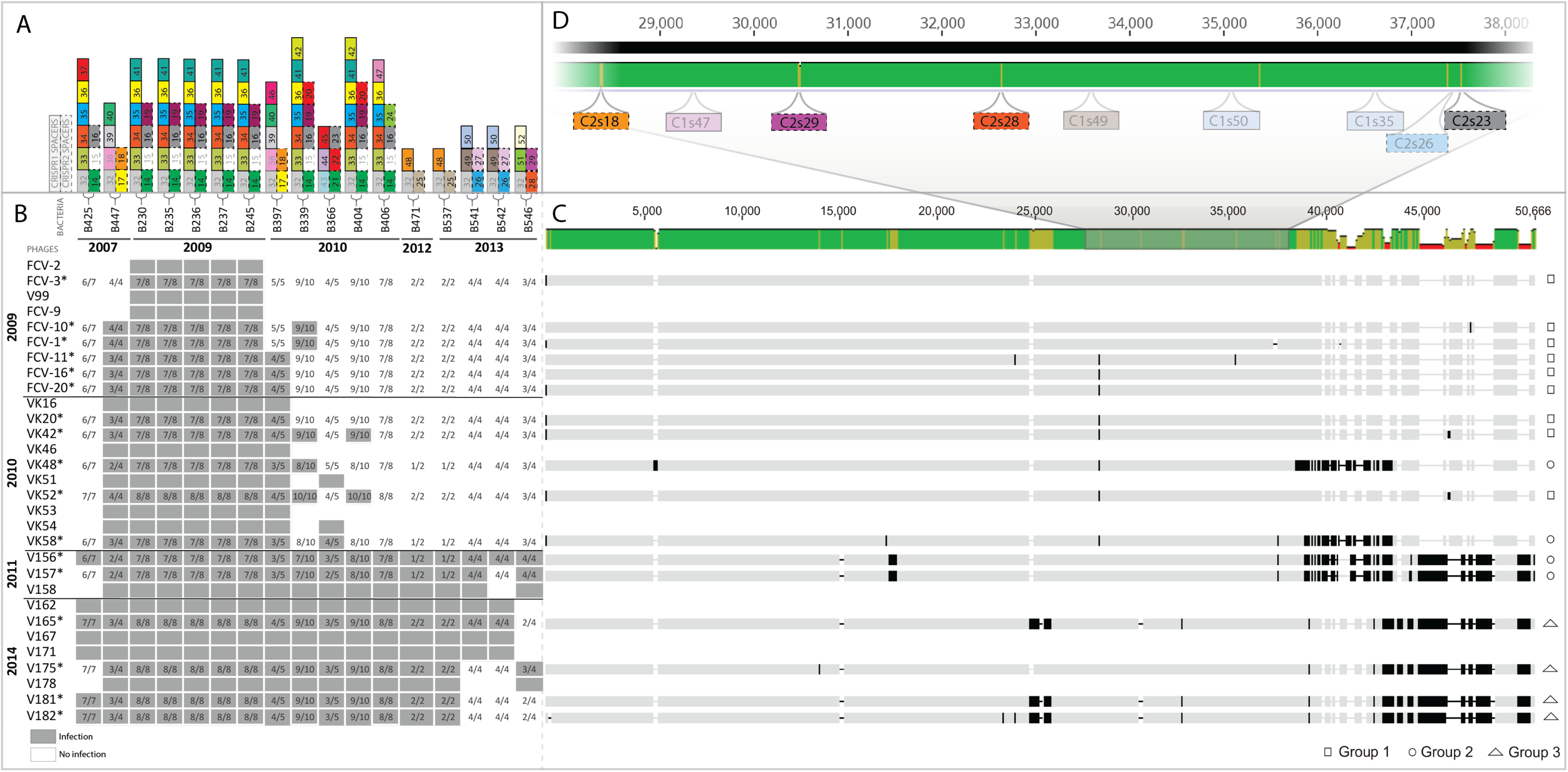
Effect of long-term arms race coevolution on phage infectivity and genome, and on host bacterial CRISPR content. **A)** Spacer diversity in the variable ends of both CRISPR loci (C1 and C2) is indicated by numbered and coloured rectangles above each bacterial strain (each number and colour referring to a specific spacer within a locus). C1 spacers are marked with solid lines (left spacer column) and C2 spacers with dotted lines (right spacer column). Numbers and colours are locus-specific and only variable spacers are shown. **B)** Host range of phages (rows) infecting *Flavobacterium columnare* isolates (columns). Gray rectangles indicate infection (presence of plaques) and white rectangles indicate resistance. Numbers within the infection/resistance rectangles indicate the overall number of CRISPR spacers with an identical match in the phage genome. **C)** Patterns of molecular evolution in the phage genomes in response to host range. Each sequenced phage genome follows the corresponding host range row from panel B. Green color in the consensus sequence (above) indicates identical sequence, yellow 30% and red < 30% identity. Black indicates nucleotide differences and insertion elements. **D)** Close-up: CRISPR protospacer regions mapped to a ~10 kb portion of the multiple sequence alignment, where the four highlighted spacers indicate phage protospacer mutations.

In multiple sequence alignment, the phage genomes clustered in three major groups (Figure 2). Group 1 phages (7 phages from 2007-2009) were identical. In the Group 2 phages (4 phages from 2010-2011), genomic changes were located at 38 922-43 395 bp (consensus sequence, in phage VK48 starting from 38 421 bp), including the absence of a predicted HNH-endonuclease. However, the nucleotide differences in a putative helicase-primase were not reflected at the amino acid level nor in the phage host range. Together with the surrounding hypothetical proteins, this genomic region might be associated with phage replication. Furthermore, these differences were missing in the Group 1 and Group 3 phages (supplementary information).

The broadest host range phages (Group 3 with 4 phages; isolated in 2014) had changes in putative structural proteins (ORFS 36 and 37, see above). Interestingly, they, as well as two phages isolated in 2011 from the Group 2 had more DNA at the terminal end of their genomes, coding additional predicted ORFs. All these additional ORFs locate upstream of ORF66, which has a putative ICE (Integrative and Conjugative Element) domain. Together with changes in the bacterial CRISPR spacer content (see below), these genomic differences seem to correlate with a broader phage host range, suggesting a mechanism necessary to overcome the bacterial constitutive defence mechanism. The additional ORFs were assigned as hypothetical proteins. One of these ORFs (no. 67.1) has a conserved domain belonging to the family of N-acyltransferases (Supplementary Figure S2, Supplementary Table S2). Another ORF (no 74.1) is a putative YopX protein, having also a mobile and extrachromosomal element and prophage functions.

In addition, phage infectivity was significantly different in the three genomic groups (groups 1, 2 and 3), interpreted as number of plaques produced over all bacterial hosts (χ^2^ = 77.252, df =2, p< 0.001, Supplementary Figure S2).

### Characterization of the CRISPR loci and PAM sequences

Next, using a previously published genome sequence (22) and our own NGS data, we identified two CRISPR loci found in all 17 *F. columnare* isolates used in this study, designated here as C1 (CRISPR1, ATCC49512: 391567-394479) and C2 (CRISPR2, ATCC49512: 1680008-1680571) (Figure 3, Supplementary Table S5). C1 contains Cas9 followed by a repeat-spacer array and two smaller cas genes (Cas1 and Cas2) in opposing directions (Figure 3). Based on cas gene composition^15^ and Cas9 homology^30^, C1 is a type II-C CRISPR locus. C2 represents the recently discovered type VI-B CRISPR system with its large RNA-targeting single-component CRISPR endonuclease Cas13b. Csx27 and Csx28, small cas gene regulators associated with some type VI-B systems, are missing from *F. columnare* as noted by^31^. We did, however, discover two ORFs downstream of the C2 repeat-spacer array whose translations show possible transposase motifs. These small, nearly identical ORFs may be remnants from possible horizontal transfer of this locus, see e.g.^32^. All 18 C1 spacers are derived equally from both phage strands and from both coding and non-coding regions. Interestingly, all 15 C2 spacers are targeting predicted ORFs on the phage coding strand. This supports the notion that type VI-B systems are targeting viral transcripts^31^.

**Figure 3.**
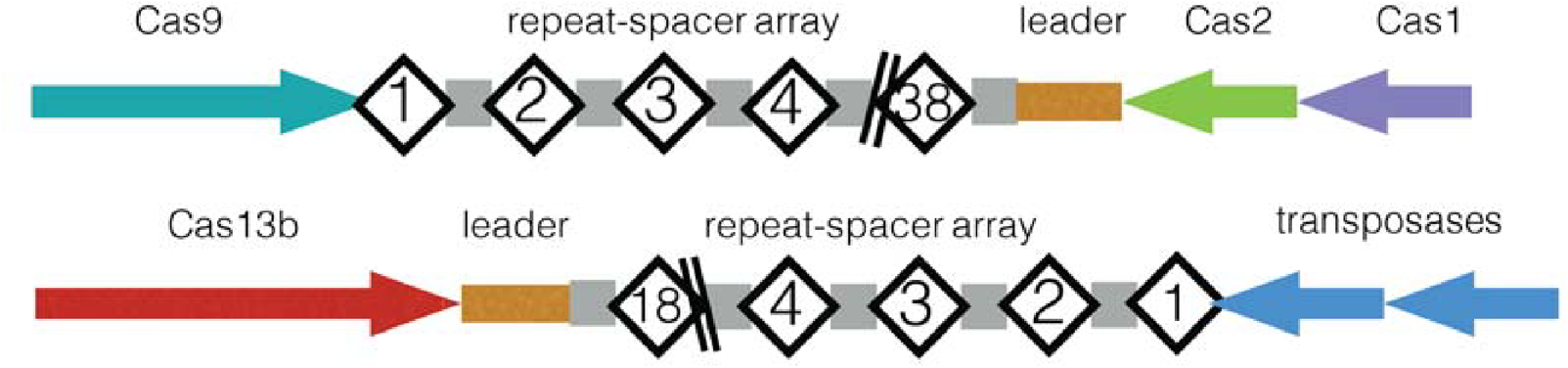
Organization of C1 and C2 loci in *F. columnare*. Spacers are numbered starting from the conserved end. Repeats are depicted by grey rectangles.

We searched for PAMs by analysing 20 bp regions surrounding each of the 18 C1 protospacers using the guide-centric approach^33^. Alignment of the results with WebLogo revealed a putative 3’ PAM with the sequence NNNNNTAAAA (Figure 4, Supplementary Table S6). This is the longest PAM sequence described to date (10 bp) if the 5 bp linker sequence is included^33^. The *F. columnare* C1 PAM was briefly addressed in a previous study^34^, in which the authors report a similar yet truncated version of the sequence described here.

**Figure 4.**
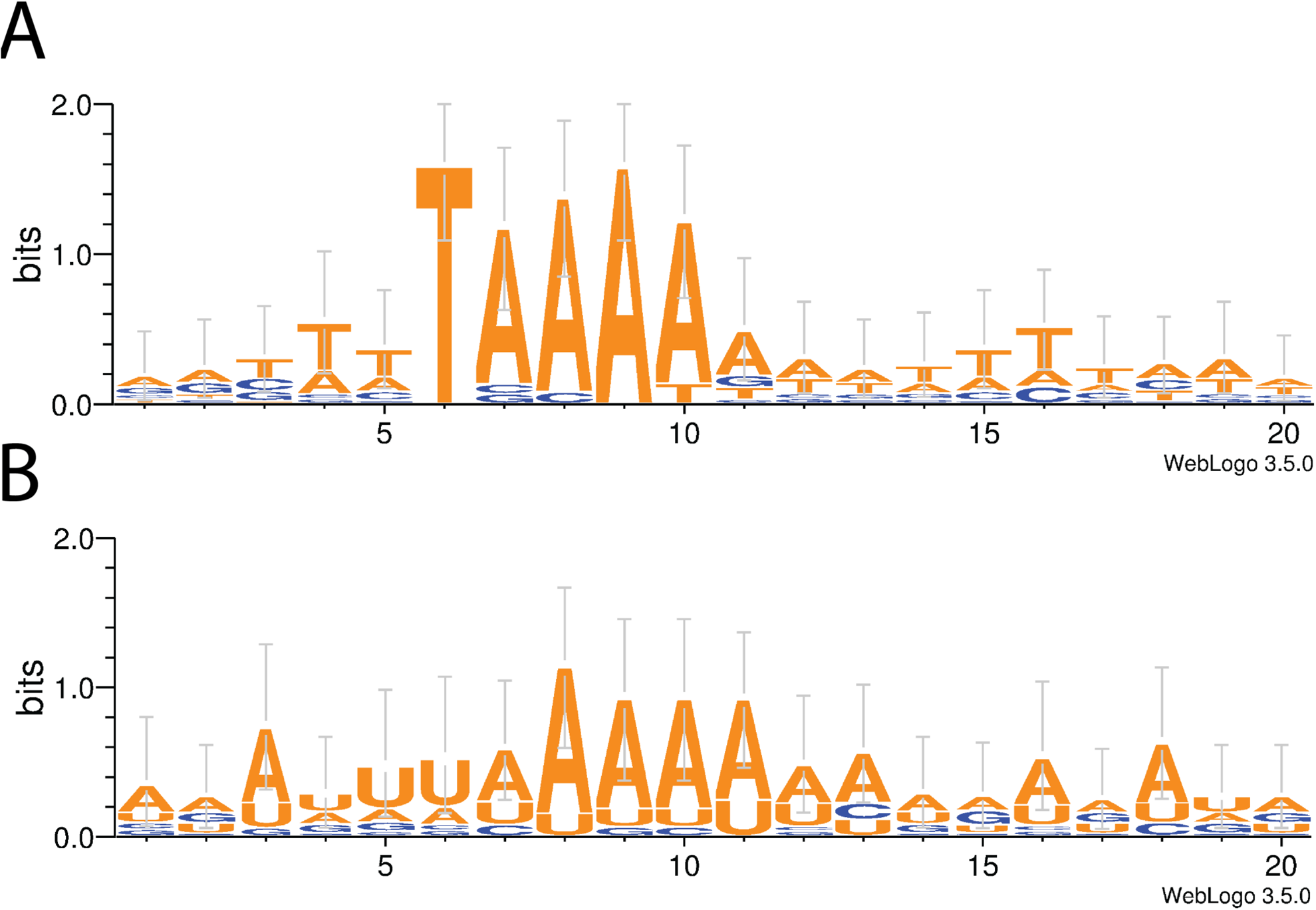
Weblogo analysis of 20 bp sequences downstream of A) CRISPR1, B) CRISPR2 protospacers in *Flavobacterium columnare* phages.

Analysing protospacer flanking sites (PFSs) associated with the 15 C2 RNA-protospacers revealed a preference for U or A in all but one 5’ PFSs (Supplementary Table S7). Also, in the 5’ PFS of spacer C2s18, A is replaced by C in some phage genomes. These findings are in line with previous studies^31^ showing that in type VI-B systems a 5’ PFS of A and U enhance interference, while C acts as an inhibitor. In the case of 3’ PFSs, either or both of the previously reported patterns (NAN or NNA) were found in 10 of the 15 C2 protospacers. Surprisingly, we also found similarities with the wider 3’ region of C2 and the 3’ PAM sequence of C1, suggesting possible interplay between these two loci in spacer acquisition (Supplementary Discussion).

### Bacterial CRISPR profiling reveals specificity of the phage-bacterium coevolution

Over time, all bacterial strains displayed fluctuating phage-matching spacer content while maintaining a set of conserved spacers. Whereas most of the conserved spacers between the strains were targeting unknown sequences or the bacterial genome itself, the variable ends of the arrays consisted almost solely of phage-targeting spacers (Figure 2, Supplementary Tables S8 and S9), exemplifying how exclusively CRISPR evolution is driven by the phages in close proximity in time and space. Furthermore, individual bacteria contained several spacers matching individual phages (Supplementary Figure S3), demonstrating the specificity of the host-parasite interaction.

On multiple occasions, the introduction of novel spacers in the bacterial population correlated with the appearance of altered protospacer sequences in phages isolated at a later time point. This was most prominent on a roughly 11 kbp area (27 352 – 38 174 in the consensus sequence), which is predicted to contain genes involved in phage replication and is being targeted by nine spacers (Figure 2, Supplementary Table S9). Across this area, six mutational sites (1 to 12 bp in length) appear, four of which are found on protospacers and whose appearance correlates chronologically with the introduction of the corresponding C2-spacers to the bacterial population (Supplementary Table S10). Furthermore, the absence of a spacer in later bacterial isolates was often followed by the reappearance of the original protospacer sequence in later phage isolates, suggesting relaxed selection towards these phages.

## Discussion

Our results highlight the long-term arms race coevolution of hosts and parasites in the aquaculture environment down to the molecular level. In accordance with previous findings^1,8-10^, the bacterial population was generally resistant against phages from past time points and susceptible to infections by (broad host range) phages from the future. Further molecular characterization pinpointed genetic changes in both host and phage populations that fit the assumption of arms race coevolution. This allows estimations of the role of constitutive and adaptive immunity as drivers of parasite infectivity in environmental populations. These results are especially important for economically relevant conditions that might be interesting for phage therapy approaches.

The evolution of generalist parasites has been shown to arise in response to host resistance in experimental settings^1,35^. Combined with phenotypic resistance patterns, our data suggest that the evolution of host immunity caused directional selection on phage infectivity and host range via genomic expansion (Figure 2, Supplementary Figure S2). In correlation with higher infectivity (Supplementary Figure S1), the genomes of the most recently isolated phages had larger genome size, increasing from 46 448 bp (in 2009) to 49 121 bp (in 2014). All additional DNA was concentrated in the terminal regions of the genomes. Although no clear functions for the three additional ORFs were predicted, structural similarity to a YopX-like protein was found. This protein has been suggested to be linked with the phage life cycle^36^, and together with the other additional ORFs (and deletion of less efficient ones) could provide improved replication and/or host lysis. Indeed, when studied over all bacterial isolates, the host generalist phages isolated most recently were most infective (Supplementary Figure S1). This likely results from a combination of phage characteristics. From an evolutionary perspective, parasite host range is expected to correlate negatively with fitness^37^. Our finding of parallel increase in host range and infectivity demonstrates that the evolutionary trajectories of the phage-bacterium interactions are not yet fully understood.

Genome and gene family expansions have been shown to be the starting points of evolutionary innovations in eukaryotic parasites^38^. These gene family expansions are often located in the recombinogenic parts of the genome, which could also apply to the phage genomes in our dataset, as the additional ORFs are found upstream of ORF66, which contains a putative ICE (Integrative and Conjugative Element) domain. ICEs integrate in chromosomes and mediate horizontal gene transfer in bacteria^39^, but their possible functionality in viruses has not been characterized. Nevertheless, these phage genome expansions demonstrate the role of the surrounding environment as a source of novel DNA and highlight the importance of long-term environmental sampling in understanding phage evolution. Yet, as the physical characteristics of the capsid strongly constrain phage genome size, our data raises new questions on what limits the phage host range, virulence and genome evolution, and what the associated trade-offs are.

The presence of CRISPR/Cas systems has been found in approximately 40% of studied bacterial species^32^, and the molecular mechanisms of different CRISPR systems are under rigorous research^40^. Some repeat-spacer arrays remain relatively constant over millennia^41^, while in some cases no two identical CRISPR loci are found in cells living in close proximity^22^. A study on a closely related species, the fish pathogen *Flavobacterium psychrophilum*, identified only inactive CRISPR loci across multiple strains^42^. In *F. columnare* we report two dynamic CRISPR loci: C1, a type II-C system associated with a novel PAM and C2, a recently identified type VI-B system. Both of the loci accumulate novel (phage-originating) spacers extending the backbone of conserved spacers. On several occasions, the introduction of CRISPR spacers to the bacterial population led to the appearance of phage isolates with specific modifications in the corresponding protospacer sequences. This is consistent with studies demonstrating the appearance of CRISPR-escape mutants due to altered protospacers^6,18^ or PAM regions^43^. While we cannot exclude the possibility that the genetic patterns of CRISPR spacers and phage protospacer regions and arms race-like counter-adaptations to putative receptor mutations (suggested by changes in ORFS 27, 28, 36 and 37 coding for phage structural proteins) observed in this study are partially the result of random drift, the conditions for host-driven selection in a natural population seem favourable.

Previous studies have addressed spacer acquisition bias in DNA-targeting CRISPR systems^5^, but studies on type VI spacer bias are missing. Unravelling patterns in target-preference may yield crucial information on the role of specific genes for viral proliferation, since RNA-targeting enables the CRISPR system to degrade viral transcripts as they are expressed in the host. Although C2 protospacers are scattered around the phage genomes, they are mostly concentrated towards the terminal regions of the genomes. In our data, target-ORFs which exhibit variability in their protospacers over time (suggesting CRISPR-evasion) include (among other ORFs with no predicted function) a predicted helicase and an exonuclease (Supplementary Table S3). Targeting these genes involved in phage replication and DNA-processing might therefore present one favourable strategy for a bacterium under phage invasion. A more detailed view of the spacer bias would, however, require the deep-sequencing of a coevolving experimental host-pathogen population.

The bacterial and phage isolates used in this study originate from a fish farming environment, a semi-natural system with a constant flow of incoming environmental water. It is possible that although attempts to isolate *F. columnare* phages from waters outside fish farming have been mainly unsuccessful^28^, phages and bacteria may have already interacted in nature, and the farming system enriches the phage population sizes to detectable levels. However, as a man-made environment, the phage-bacterium interaction is subjected to selection by farming practices, especially high fish densities and the use of medication, which may select for the most virulent and antibiotic tolerant *F. columnare* strains^44^. Thus, by providing essential information on the phage-bacterium interactions under multifactorial selection in these settings, our data and study system are directly relevant for sustainable use of phage therapy applications in field conditions. For example, in laboratory conditions eliciting phage resistance in *F. columnare* has been shown to cause reversible morphotypic changes that can maintain constitutive resistance^45,46^. These resistant morphotypes are, however, missing from natural isolates, probably because of their impaired growth, tolerance to protozoan predation and capacity to infect fish^45,47^. This indicates higher benefits of CRISPR immunity under multifactorial selection in field conditions, although our data show that both constitutive and adaptive bacterial resistance mechanisms are active and important in natural settings.

CRISPRs could not completely explain the resistance patterns of the host, possibly due to high phage pressure in the experimental setup^48^. However, under low phage selection (such as natural aquatic settings) CRISPRs may still be preferred over mutation-based immunity due to their minimal effects on overall fitness of the host^49^ at least when not constantly expressed^50^. Consistent with a previous suggestion^51^, the presence of the high diversity of CRISPR spacers (e.g. in 2010) could have reduced phages’ ability to overcome bacterial resistance by point mutations, and may have caused larger shifts in the structure of the phage population, observed here as phages with increased genome size. Considering the costs related to maintaining broad resistance against phages, combining constitutive immunity and diverse CRISPR immunity may be the most successful strategy for a natural bacterial population^52^, as the phage have the potential to rapidly overcome these bacterial defence mechanisms to ensure their own persistence. The expanding host range driven by the increased genome size is likely to be restricted by the physical capacity of the capsid. Nevertheless, other factors that limit the evolutionary potential of this specific phage-bacterium interaction have yet to be discovered. Furthermore, the expansion of the host range may trade-off with phage infectivity^37^, which might be the point at which the adaptive CRISPR defence alone would be sufficient to overcome those infections.

The importance and volume of aquaculture is growing steadily to meet the increasing need for high-quality protein for human consumption^53^, and new methods, such as phage therapy, to control and manage bacterial diseases are under rigorous research^54,55^. The greatest claim against phage therapy is the evolution of resistance in the target bacteria, making phages a “once only” solution. Our data on the long-term phage-bacterium coevolutionary dynamics demonstrate the plasticity of phage infectivity and their capacity in overcoming bacterial resistance mechanisms in settings relevant for phage therapy and can help in estimating resistance dynamics in real-life scenarios.

## Materials and methods

### Phage and bacterial isolates used in this study

Phage and bacteria were isolated from a private fish farm (from fish and tank water) in Central Finland and from its immediate surroundings from inlet and outlet water (Supplementary Tables S1 and S2). Bacterial strains were isolated from water samples by plating on Shieh agar^56^ or Shieh agar supplemented with tobramycin^57^, as previously described^58^. Among the isolates, bacteria belonging to genetic group C were chosen based on ARISA genotyping using the methodology described previously^58,59^. The interaction between *F. columnare* and its phages is genotype-specific, which allowed us to monitor the evolution of phage-bacterium relationship in this study using a specific host genotype. For this study we used all available bacterial strains isolated from the fish farm and its surroundings that belong to the genetic group C. Bacterial isolates were stored frozen at −80°C, with 10% Fetal Calf Serum and 10% glycerol.

Phages were isolated from water samples by enrichment. A filtered water sample (pore size 0,45 μm, Nalgene) was used to dilute five-fold Shieh medium, and an overnight-grown enrichment host (belonging to genetic group C) was inoculated in this medium. Cultures were grown at room temperature (23 °C), at 110 rpm on a benchtop shaker (New Brunswick Scientific) until they turned turbid. A double agar layer method was used for detecting phages: 300 µl of turbid sample with 3ml of 0,7% soft Shieh agar (tempered to 47 °C) was applied on solid Shieh agar. After incubation (23 °C) for 24-48 hours (depending on the bacterial growth) plaques were picked. Three rounds of plaque purification were performed for each phage isolate. For further characterization, phage lysates were prepared adding 5 ml of culture media on a plate with confluent lysis, and shaken at 8 °C, at 95 rpm on a benchtop shaker for 6 hours. Phage isolates were stored frozen in −80°C with 20 % glycerol. Phages were characterized by genome restriction profiles using prior EcoRI experiments (data not shown), and for this study we used all the phage isolates infecting the bacterial genetic group C.

### Phage host range

For analyzing phage infectivity and host range, an overnight-grown bacterial culture (in Shieh medium, 23°C, 110 rpm) was plated using the double agar overlay method by mixing bacterial culture 1:10 with 0.7% soft Shieh agar tempered to 47 °C and pouring this on agar plates. The phage was diluted until million fold (10^−6^), and 10 µl samples of each dilution were spotted on top of the bacterial lawn. All bacterial strains were infected with all phages in a pairwise manner. After incubation of 48 h, the plates were checked for single plaques. Plaque formation was considered as a positive result for phage infectivity.

### Time-shift experiment and statistical analysis

Bacterial resistance in response to phages from past, contemporary and future time points was analysed with generalized linear mixed models using SPSS 22.0. The response variable of phage infection was encoded as a binary trait, therefore binomial distribution with logit link transformation was used. In the analyses time-shift was used as a fixed factor and phage isolation year as a random factor with bacterial identity as a blocking factor (subject). Only phages able to produce plaques in the host bacteria were considered infective in the analysis.

### Phage genome sequencing

Phages subjected to genome sequencing were chosen based on their infection profiles (Figure 2). Phage DNA was isolated using a protocol described by Santos^60^ with slight modifications. Briefly, after RNase (10 μg/ml) and DNase (1 μg/ml) treatment phages were precipitated with 40 mM ZnCl_2_ and incubated for 5 minutes followed by pelleting (15 000 × *g*, 5 min) and resuspended in TES-buffer (0.1 M Tris-HCl, pH 8; 0.1 M EDTA; 0.3% SDS). DNA was purified using a GeneJET^TM^ Genomic DNA isolation kit column (Fermentas). 100 ng of DNA was subjected to genome sequencing with Illumina MiSeq in Institute for Molecular Medicine Finland (FIMM), apart from isolate FCV-1 which was sequenced using Roche 454 at LGC Genomics, Germany.

### Genome assembly

The phage genomes were assembled using Velvet-assembler (v. 1.2.10)^61^ with optimal k-mers per genome (average coverage ~1600x). Differences appearing in only one genome were checked and corrected if needed. Single insertions and nucleotide differences were checked by sequencing directly from the genome using specific primers (Supplementary Data Table S10). Open reading frames (ORF’s) were predicted using GeneMarkS (v. 4.28)^62^ and Glimmer (v. 3.02b)^63^. Protein Homology/analogY Recognition Engine V 2.0 (PHYRE^2^)^64^ was used for predicting protein structures.

### CRISPR sequencing and analyses

Bacterial DNA was extracted from overnight-grown turbid cultures using the GeneJET^TM^ Genomic DNA isolation kit (Fermentas). We used the previously published *Flavobacterium columnare* ATCC 49512 genome^65^ and our own unpublished NGS data as references in designing primers for amplifying the CRISPR loci. NetPrimer (Premier Biosoft) was used for the analysis. Primers (Supplementary Data Table S10) were provided by Sigma. CRISPR loci from the extracted DNA were amplified in a PCR reaction (PIKO Thermal cycler by Finnzymes) using the Phusion Flash II polymerase in a 20 µl reaction with a template concentration of 0.5 ng/µl. The protocol was completed following the manufacturer’s instructions. The annealing temperature was 65 °C for CRISPR1 and 67 °C for CRISPR2, and the elongation step was 30 sec for CRISPR1 and 27 sec for CRISPR2. The reactions were purified for sequencing using the QIAquick PCR Purification Kit (Qiagen). The amplified PCR products were sequenced with the Sanger method. Raw data were transformed using SequenceAnalysis 6 basecalling. Two replicates were done for each read and the consensus sequences were manually determined using Geneious 9.1.4 (Biomatters Ltd).

The determination of the orientations of the CRISPR loci (namely, the transcription direction of the repeat-spacer arrays) was based on analyses using CRISPRDetect^66^, comparisons with other similar CRISPR systems and observations the polarization of spacer acquisition. Unlike in most CRISPR systems, the crRNA in type II-C systems is transcribed starting from the conserved end^67^. By assuming this transcription direction, we could detect a conserved PAM sequence downstream of C1 protospacers. As all type II systems described thus far exhibit downstream PAMs, this reinforces the assumption that the C1 repeat-spacer array is also transcribed starting from the conserved end. Very low confidence values associated with C1 direction prediction by CRISPRDetect led us to not use this tool with this locus. However, CRISPRDetect was able to produce confident direction prediction values for the C2 repeat-spacer arrays, which are in line with those of a previous study^31^ and which place the transcription initiation point at the variable end, as in most CRISPR systems. This direction also enables the spacers to target mRNA from predicted viral ORFs and produces PFSs in line with those of a previous study^31^. Spacers were numbered starting from the conserved end, so that each spacer in the total spacer pool had a unique sequence ID (e.g. C1s13). In C1, a 114 bp area containing degenerate repeats between spacers 4 and 5 was skipped, and numbering was resumed downstream of this area.

### Analysis

Geneious (v. 9.1.4) was used for the annotation and analysis of the CRISPR sequence data and phage PAM and PFS extractions. WebLogo (v. 2.8.2) was used to create sequence logos of the PAM and PFS sequences.

## Acknowledgements

This work was supported by the Finnish Centre of Excellence Program of the Academy of Finland; the CoE in Biological Interactions 2012-2017 (#252411), by the Academy of Finland grant #266879, and by the Jane and Aatos Erkko Foundation. The authors would like to thank MSc Jenni Marjakangas, MSC Katja Ryymin and Mr. Petri Papponen for skillful help in the laboratory, Dr. Heidi Kunttu for kindly donating genotyped bacterial strains, and Dr. Reetta Penttinen for help in bacterial isolation. In memory of Prof Jaana Bamford.

